# Parallel Analyses by Mass Spectrometry (MS) and Reverse Phase Protein Array (RPPA) Reveal Complementary Proteomic Profiles in Triple-Negative Breast Cancer (TNBC) Patient Tissues and Cell Cultures

**DOI:** 10.1101/2024.05.30.596640

**Authors:** Nan Wang, Yiying Zhu, Lianshui Wang, Wenshuang Dai, Taobo Hu, Zhentao Song, Xia Li, Qi Zhang, Jianfei Ma, Qianghua Xia, Jin Li, Yiqiang Liu, Mengping Long, Zhiyong Ding

## Abstract

High-plex proteomic technologies have made substantial contributions to mechanism studies and biomarker discovery in complex diseases, particularly cancer. Despite technological advancements, inherent limitations in individual proteomic approaches persist, impeding the achievement of comprehensive quantitative insights into the proteome. In this study, we employed two widely used proteomic technologies, Mass Spectrometry (MS) and Reverse Phase Protein Array (RPPA) to analyze identical samples, aiming to systematically assess the outcomes and performance of the different technologies. Additionally, we sought to establish an integrated workflow by combining these two proteomic approaches to augment the coverage of protein targets for discovery purposes. We used fresh frozen tissue samples from triple-negative breast cancer (TNBC) and cell line samples to evaluate both technologies and implement this dual-proteomic strategy. Using a single-step protein denaturation and extraction protocol, protein samples were subjected to reverse phase chromatography (LC) followed by electrospray ionization (ESI)-mediated MS/MS for proteomic profiling. Concurrently, identical sample aliquots were analyzed by RPPA for profiling of over 300 proteins and phosphoproteins that are in key signaling pathways or druggable targets in cancer. Both proteomic methods demonstrated the expected ability to differentiate samples by groups, revealing distinct proteomic patterns under various experimental conditions, albeit with minimal overlap in identified targets. Mechanism-based analysis uncovered divergent biological processes identified with the two proteomic technologies, capitalizing on their complementary exploratory potential.

## Introduction

Proteins are functional biomolecules to directly dictate almost every biological process and thus are being the focus of biomarker discovery in translational medicine [1, 2]. Quantitative proteomics, especially high-plex proteomic technologies have become the front-line analytical approach in discovering novel protein biomarkers in many disease settings [3, 4]. Amongst those, mass spectrometry (MS)-based methods have emerged as a key strategy of choice for qualitative and quantitative detection of proteins in biological samples. In the preceding decade, MS has become the dominant player in this field, owing to its discovery capability theoretically without a prior knowledge of the analytes. Nowadays, analytical methods utilizing high performance liquid chromatography (HPLC) for peptide separation coupled with electrospray ionization (ESI) and tandem MS/MS spectrum-based detection widely adopted for global protein identification, while quantification could be either label-based or label-free methods [5]. Efforts have also been made to improve the detection sensitivity and quantification accuracy and those are mainly focused on three technical dimensions. 1. To increase the protein detectability by optimizing the protein recovery and peptide digestion efficiency in sample preparation prior to liquid chromatography. 2. To incorporate orthogonal separation techniques including the ion mobility technology in the front-end of the mass spectrometer. improve the analytical robustness of the front-end mass analyzer. 3. To advance mass spectrum scanning modes and data analysis software to improve proteome coverage per unit time and robustness of protein quantification. [6, 7]. While MS-based discovery proteomics hold central promises in biomarker profiling, other techniques also exist and particularly in the field of cancer biology, reverse phase protein array (RPPA) has been highlighted as an excellent experimental approach for cancer-related biomarker profiling [8, 9]. As originally designed to address signaling pathway alteration in cancer, RPPA is tactically suited for targeted proteomics in a high-throughput format. Due to the highly sensitive antibody-based detection and amplification method, RPPA focuses on proteins in key signaling pathways, transcription regulation and their modified proteoforms that are low protein expressors but major functional determinants in pharmacodynamics and druggable targets during oncogenic processes[10–12]. Nevertheless, RPPA is confined by the availability of high-quality antibodies as well as its semi-automated workflow and in a typical experimental assay, panelized detectable targets ranging from 50 up to 450 [13–15].

Clinical tissue samples bearing invaluable biological information are most commonly used materials to address oncogenic questions such like the molecular alterations arising from cancerous origins, different progression stages, primary and metastatic sites or treatment/response associated phenotypic effect. Amongst a wide array of sample types, fresh frozen (FF) tissues are still the preferred material of choice for high-plex discovery proteomics due to its biological integrity preserved and therefore are widely used in tissue-based proteomic profiling such as MS or RPPA [16–19]. In MS experiments, sample preparation is an essential part in proteomic characterization of clinical tissue samples. Lysing and extraction of proteins from clinical samples require different organic solvents and detergents with additional tissue disruption and homogenization processes including sonication and physical disruption [20]. Although, organic solvent-based extractions (such as 2,2,2-Trifluoroethanol-based method) were reported in application of FF-based proteogenomic discovery, traditional detergent-based extraction methods have long been regarded as the gold-standard approach in tissue proteomics setting [21, 22]. Denaturants including ionic and non-ionic agents such as urea, guanidine HCl, SDS, Triton X-100 and NP-40 are efficient lysing reagents to disrupt tissue and solubilize protein complexes including membrane proteins. Those detergents are typically removed using different purification techniques to allow more efficient digestion and prevent adverse chemical deposition in MS instruments[23, 24]. FF tissues are also favored clinical resources in RPPA experimental settings and this was proven in large-scale pan-cancer multi-omics profiling primarily featured by the cancer genome atlas (TCGA) project. FF sample preservation processes developed for RPPA were described. Currently, generally accepted methods use denaturing agents such as urea, thiourea, SDS and Triton X-100 all of which have shown decent compatibility with FF samples [13, 25, 26]. Of further technical interest, there were also studies reporting on a single lysing procedure compatible for both MS and RPPA mediated protein profiling, however, this work assessed the quantitative MS profiling using difference in gel electrophoresis (DIGE) for protein separation/selection and followed by matrix absorbed laser dissociation ionization mass spectrometry (MALDI-MS) rather than liquid chromatography-based MS (LC-MS)[27].

In addition, from cancer biomarker discovery perspective, bioinformatic analysis on cell line models testing the predictive power for drug sensitive evaluation revealed the added benefit of incorporating both MS and RPPA strategies in biomarker profiling. In this study, using GI50 values as responsive variables, multi-omics data at genomic, transcriptomic and proteomic levels derived from NCI-60 cell lines with drug response measures of 47 FDA-approved cytotoxic and targeted agents were evaluated [28]. Main conclusions have been drawn where it showed significantly increased predictive power by inclusion of both MS and RPPA data and both datasets provided complementary information contributing towards the response prediction[28]. These findings point at the potential of cross-platform application in aid of protein biomarker discovery and mechanistic elucidation, however apart from the abovementioned single lysis solution used in tissue protein extraction followed by RPPA and DIGE workflow, there is yet no such experimental approach established compatible for both LC-MS and RPPA.

Proteomic landscapes in multiple cancers have been established systematically through RPPA (TCGA) and MS (clinical proteomic tumor analysis consortium CPTAC). In breast cancer (BC), comprehensive proteogenomic profiling has led to deeper understanding of proteomic driven molecular alteration linking to their genetic traits and plausible therapeutic targets [19, 29]. In the current study, we used primary both FF tissues from triple negative breast cancer (TNBC) as well as in vitro cell lines (293T, MKN7 and OVCAR3) as models to assess an in-house developed all-in-one workflow for joint high-plex proteomic profiling using both LC-MS and RPPA and compared their analytical strength and weakness in addressing biological questions respectively. This work set as a preliminary work that may be shared to the broad community for mechanistic exploration and biomarker profiling.

## Materials and methods

### Clinical sample acquisition and characteristics

The study was approved by the Peking University Cancer Hospital ethics committee (reference number 2020KT113). Tumor and paired normal tissues were obtained from surgical specimens of seven patients at Peking University Cancer Hospital. These patients were diagnosed by core needle biopsy and had not received any systematic therapy before surgery. The samples were obtained immediately after the surgery by a pathologist. The paired normal tissue was obtained at least 2cm away from the tumor margin. The detailed clinicopathological information of the included patients is listed in Supplementary Table 1.

### Lysis buffer composition

The lysis buffer was prepared to contain the following components: 50 mM HEPES (pH 7.4), 150 mM NaCl, 1 mM EGTA, 10 mM sodium pyrophosphate, 100 mM sodium fluoride, 1.5 mM MgCl2, 1% Triton X-100, 10% glycerol, and 1 mM sodium orthovanadate. Additionally, proteinase inhibitor (Roche 05056489001) and phosphatase inhibitor (Roche 04906837001) were added following the manufacturer’s instructions. The resulting lysis buffer was stored at –20□ and thawed on ice prior to use.

### Clinical sample preparation and protein quantification for dual proteomic profiling by LC-MS and RPPA

Fresh frozen TNBC tissues and paired normal tissues were semi-thawed and weighted for subsequent processing in grinding tubes. For each sample, ice-cold lysis buffer was added at a ratio of 1:20 (sample/buffer). Depending on available materials, we used 45-80 mg of each sample to allow parallel LC-MS and RPPA profiling. For tissue disruption, 6 ceramic beads were added to each sample grinding tube and loaded into the pre-cold chamber (–5□) of the beads ruptor (OMIN bead ruptor 24 Elite). Tissues were disrupted with the following settings: 2 cycles, each lasting 30 seconds with a 10-second cooling time interval. Homogenized samples were then subjected to centrifugation (14,000 rpm) for 15 minutes at 4□, and the resulting supernatants were retained for protein quantification using a standard BCA protocol (Pierce BCA kit 23225) and a microplate reader (Biotek, Epoch2).

### Cell line sample preparation

Cell lines (293T, MKN7 and OVCAR3) were obtained from ATCC. All cell lines were verified by Satellite Tandem Repeat (STR) to ensure authenticity. Cells were cultured at 37□with a 5% CO2 supply. For 293T cells, serum starvation was conducted in serum-depleted DMEM medium for 24 hours before stimulation with 10% fetal bovine serum (FBS). Cells were harvested 30 minutes post-treatment. MKN7 and OVCAR3 cells were cultured in complete medium (RPMI1640 with 10-20% FBS) and harvested when cells reached over 80%-90% confluence.

## Label-free Liquid chromatography mass spectrometry (LC-MS)

### Sample purification

Solubilized protein extracts were purified by acetone precipitation. Specifically, 80 μl of precooled acetone was added to 50 μg of each sample and placed in –20□overnight. Precipitated protein was pelleted by centrifugation at 16,000 x g for 10 mins. Supernatants containing detergent and salt were discarded. Protein pellets were washed with cold acetone 3 times, then dried for 10 mins at room temperature (RT). Samples were resolubilized in a buffer made of 8M urea and 50 mM ammonium bicarbonate (AMBIC), pH 7.8, and sonicated. Protein disulfide bonds were reduced by 5 mM of dithiothreitol at 30□ for 1 hour and alkylated by freshly made 10 mM of iodoacetamide at RT for 30 mins in the dark. After urea was diluted in 4 x 50 mM AMBIC buffer, MS grade trypsin (Promega) was applied to a final protease-to-protein ratio of 1:25 (w/w). After overnight digestion at RT, samples were acidified by trifluoroacetic acid (TFA) and further purified by C18 stage tips. Each C18 tip was equilibrated by passing 50 μl of 0.1% TFA and 80% acetonitrile (ACN) solvent. Samples were then loaded to the C18 tips and washed by 55 μl of 0.1% TFA twice. Finally, peptides were eluted off the C18 tips by 10 μl of 0.1% TFA and 80% ACN solvent twice and dried in a vacuum concentrator.

### LC-MS/MS analysis

Purified samples were resuspended in 0.1% formic acid (FA), and loaded onto the autosampler of Ultimate 3000 UPLC system (Thermo Fisher Scientific). Each injection contained ∼ 250 ng of peptides, which were separated by an in-house made analytical column (30 cm, 100 µm ID, 1.9 µm C18). LC mobile phases were 0.1% FA in water as A and 0.1% FA in 80% acetonitrile as B. LC flow rate was set at 300 nL/min. For the data-dependent acquisition (DDA) method, LC gradient and MS parameters were set as follows: mobile phase B started at 4%, increased to 15% within 2 min, and then gradually raised to 37.5% within 80 mins. MS1 spectra were acquired at an MS scan range of 300-1500 m/z, an orbitrap resolution at 60K (200 m/z), a maximum injection time at 30 ms, and a normalized AGC target at 250%. MS2 spectra were collected at a scan range of 200-1400 m/z, an isolation window of 1.6, an orbitrap resolution at 15K (200m/z), a normalized HCD energy at 27 %, a maximum injection time of 22 ms, a normalized AGC target at 100%. For data-independent acquisition (DIA), parameters were set as follows: on the LC, Mobile B started at 8% and increased to 37% within 120 min. The full MS was carried out in the Orbitrap by scanning m/z 350-1,150 at a resolution of 120K (200 m/z), with an AGC target at 1E6 and a maximum injection time at 50 ms. One full MS event followed by 30 MS/MS windows in one cycle. The precursors were fragmented by HCD at normalized collision energy at 32%. The MS/MS was carried out in the Orbitrap by scanning m/z 200-1,600 at a resolution of 30K (m/z 200), with an AGC target at 1E6 and a max injection time of 54 ms. Two technical runs were generated for both TNBC and control samples with a randomized order to minimize the impact of LC-MS/MS system instability on the measurement. Three biological replicates were analyzed for cell samples.

### MS data analysis

Collected raw files from the DDA method were analyzed by Proteome Discoverer 2.4 software (Thermo Fisher Scientific). Searching was done by matching spectra to a UniProt homo sapiens database (downloaded 2021/03). The parameters were set as in the following: protease was defined as trypsin, maximum missed cleavage was 2, minimum peptide length was 6, max peptide length was 44, precursor mass tolerance was 10 ppm, fragment mass tolerance was 0.02 Da, dynamic modification was set as oxidation at methionine and acetylation at protein N-terminus, static modification was carbamidomethylation at cysteine. Percolator was employed and the filter parameter was set at 1 % false discovery rate (FDR) both at peptide and protein level. Label-free quantitation was based on extracted peak areas of peptides with minora feature mapper, which could match features between runs. Unique and razor peptides were used for quantification. Raw files collected by the DIA method were analyzed by Spectronaut v17 (Biognosys) against human fasta file (Uniprot, UP000005640) with the following settings: Enzyme was Trypsin/P. Two missed cleavages were allowed. Both peptide and protein false discovery rate (FDR) were set at 1%. Carbamidomethylation at Cysteine was set as a fixed modification. Acetyl at Protein N-terminus and oxidation at Methionine were defined as variable modifications. Normalization was based on the total peptide amount. Protein abundance was calculated on the top 3 most abundant peptides. For MS missing data handling, we applied NAguideR package [30].

## Reverse phase protein array

### Sample processing and RPPA workflow

Reverse phase protein array (RPPA) was conducted following this procedure: Protein lysates were pre-mixed with sample dilution buffer (50% glycerol, 4XSDS buffer with 6ml of beta-mercaptoethanol) to achieve a final concentration of 1.5 mg/ml. Concentration-normalized samples were then subjected to a 2-fold dilution in sample dilution buffer (a mixture of lysis buffer, 50% glycerol, and 4X SDS buffer with 6 ml of beta-mercaptoethanol at a ratio of 3:4:1). Five serial dilutions (1, 1/2, 1/4, 1/8, 1/16) were performed using automated liquid handling workstations (Tecan Fluent series). The prepared samples in 384-well plates (low-binding Molecular Devices) were deposited onto nitrocellulose-coated glass slides (Gracelab ONCYTE superNOVA) via a solid pin contact printer (Quanterix 2470 Arrayer). On-slide controls, including cell lines with or without treatment, and a mixture of cell lines and tonsil tissue lysate, were used for staining quality controls (QC) and quantitative QC measures. A similar experimental setup can be referenced in the literature [13]. In total, about 400 identical slides were prepared.

Each slide was then subjected to colorimetric signal quantification using one of a panel of 305 antibodies validated in-house, including 227 targeting total proteins and 78 targeting phosphoproteins or other PTMs (Supplementary S1). Slides were first blocked with Re-Blot (Millipore) at RT, followed by blocking with I-block (Fisher), and antigen retrieval with hydrogen peroxide (Fisher). The slides were then sequentially blocked with avidin, biotin, and protein block (DAKO) before undergoing primary antibody incubation for 1 hour at RT. Secondary antibodies (DAKO) against rabbit or mouse were then applied, followed by Tyramide Signal Amplification (TSA, Akoya) and DAB colorimetric visualization (DAKO). All staining processes were conducted automatically on DAKO Link 48 Autostainer (Agilent). Stained slides were scanned on a high-throughput slide scanner at a scanning resolution of 10 micron (Huron LE120), and images were used for downstream processing.

### RPPA data processing and analysis

The digital transformation of images was performed using MicroVigene software (version 5.6.0.8). The output text (txt) and image (tiff) files underwent SuperCurve fitting with the R package SuperCurve. This step aimed to generate expression data (rawlog2 files) and quality control (QC) data for each slide [31]. Correction factors (CF) were calculated to evaluate sample outliers intra– and inter-experimentally. For rawlog2 data normalization (loading adjustment), each column (antibody) was median subtracted column-wise and then each row (sample) was median subtracted row-wise. This generated a normalized log2 file, which was further squared to generate a linear dataset (Normlinear) (Supplementary S2). All these data sets were processed for downstream quantitative comparison and graphical visualization.

## Data analysis

All data generated from MS and RPPA profiling were processed in R (Version-4.1.0). Data cleaning, clustering, differential expression analysis, correlation analysis, and overlapping analysis were performed and plotted mainly using R packages (dplyr, ggplot2, pheatmap, clusterProfiler, limma, VeenDiagram). External RPPA proteomic data (batch normalized level 4 data) and corresponding metadata for TNBC were downloaded from Genomic Data Commons (GDC data portal: https://portal.gdc.cancer.gov/). TNBC MS data (QC-passed and normalized counts) and corresponding metadata were downloaded from Proteomic Data Commons (PDC data portal: https://proteomic.datacommons.cancer.gov/pdc/). All downloaded files are in Supplementary S3. For correlation analysis, log-transformed MS data were used for plotting, and for RPPA, normalized log2 (normlog2) data were used accordingly. The Wilcoxon ranked sum test was employed for the differential expression analysis of individual RPPA targets. Unless otherwise mentioned, a p-value of 0.05 was consistently used throughout the study as the threshold for statistical significance.

## Results

### Concurrent proteomic profiling of TNBC tissue samples using label-free MS and RPPA

To assess and compare the outcomes and performance of the two prevalent proteomic technologies, Mass Spectrometry (MS) and Reverse Phase Protein Array (RPPA), we first established a protein extraction workflow from patient tissue samples, enabling downstream quantitative analysis using both approaches. Firstly, we evaluated the protein yield from freshly frozen (FF) tissues using standard BCA. With starting FF materials ranging from 45-80 mg per sample, our approach resulted in a satisfactory protein yield in the range of 27-115 µg total protein per mg of tissue (Supplementary S4). About 40 µg of total protein in lysate from each sample was used for RPPA, and the rest protein for MS. For the LC-MS experiment, buffer exchange was conducted to remove salts and high concentrations of detergents that could potentially interfere with MS. Within an 80-minute LC gradient, over 3,300 proteins were identified in our label-free MS/MS run (Supplementary S5). After correcting for missing values, 2,583 proteins were retained to construct an unsupervised clustering map distinguishing TNBC from paired normal tissues in a near-perfect manner, with only one TNBC5 tumor sample not grouped in the tumor cluster (Fig. 1A). In parallel, RPPA generated a targeted proteomic profiling containing 305 total protein targets including 221 total proteins and 84 proteins targets including phosphoproteins and other PTMs (Supplementary S2). An unsupervised heatmap also effectively distinguishes tumor and paired normal tissues, again, with only TNBC5 tumor not grouped in the tumor cluster (Fig. 1B).

**Fig. 1.**
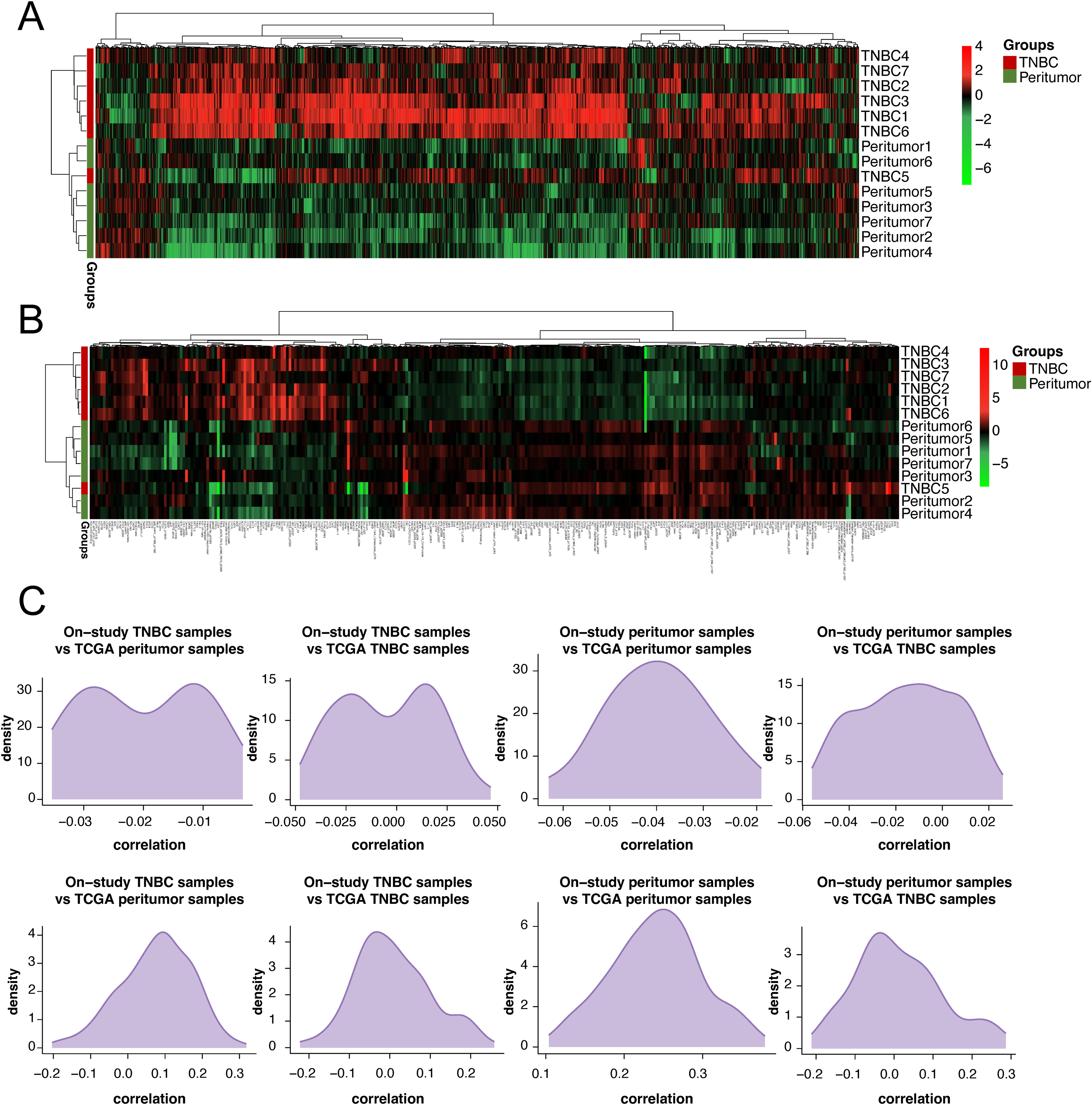
Parallel MS and RPPA proteomic profiling of TNBC and normal control tissues. A. Unsupervised hierarchical clustering of TNBC and paired normal tissues (7 versus 7) based on MS profiling. B. Unsupervised hierarchical clustering of TNBC and paired normal tissues (7 versus 7) based on RPPA profiling. C. Pairwise correlation between our MS samples and TCGA-TNBC MS samples and adjacent normal. Matched targets were used to generate sample-wise Pearson correlation and plotted using density plots. Comparisons between our on-study samples (tumor or normal) and TCGA samples (tumor or normal) were shown on separate plots. D. Pairwise correlation between our RPPA samples and TCGA-TNBC RPPA samples and adjacent normal. Matched targets were used to generate sample-wise Pearson correlation and plotted using density plots. Comparisons between our on-study samples (tumor or normal) and TCGA samples (tumor or normal) were shown on separate plots.

### RPPA data exhibits better correlation with public data than MS data

We then compared our MS and RPPA data with the public Clinical Proteomic Tumor Analysis Consortium (CPTAC) and the TCGA-TNBC RPPA data, respectively. For the public CPTAC MS data, 18 TNBC and 3 paired normal controls were obtained after filtering out QC-failed samples, with a total of 11,146 proteins identified. Of these, 2,365 proteins were matched with our identified protein (2,365/2,583) in MS and were used for subsequent correlation analysis. While our MS tumor data exhibited nearly no inter-sample correlation, it showed a slightly better correlation with matched CPTAC tumor data (Pearson R2 ranging between –0.05 and 0.05) compared to our tumor data with public normal controls (R2 between –0.03 and –0.01). When comparing our normal controls with public controls or tumors, no discernible correlations or differences in correlations were observed (Fig. 1C).

As for the RPPA data, 247 out of 305 proteins identified in our RPPA were matched to the public TCGA RPPA data (Supplementary file S3). In contrast to the MS data, a general trend of positive correlation was observed, with inter-normal comparison showing a better correlation (Pearson R^2^ ranging between 0.1 and 0.4) than inter-tumor (Pearson R^2^ ranging between 0.2 and 0.25). Our tumor data did not exhibit a correlation with either the public tumor or normal data, likely attributed to the heterogeneity of patient samples and the limited number of samples in the studies (Fig. 1D).

### MS and RPPA identifies distinct sets of differentially expressed proteins in TNBC vs peritumor tissues

Since MS and RPPA identified different sets of proteins while both proteomics profiles distinguished TNBC from paired peritumor tissues effectively (Fig. 1C), we investigated the differentially expressed (DE) proteins identified by each technology, seeking to elucidate shared biological mechanisms and those exclusively identified by either MS or RPPA. Given the broader protein coverage of the MS profiling, with 2,633 quantified targets compared to the 305 proteins in RPPA, log fold-change thresholds were set to 1.5 for MS and 0.8 for RPPA to identify DE proteins at a similar percentage. As a result, 773 (29.6% of 2633) and 84 (29.2% of 305) DE proteins were identified from MS and RPPA, respectively (Supplementary S6). Volcano plots illustrated differential expression patterns for MS and RPPA, highlighting top-ranked –log adjusted p-values (Fig. 2A/B).

**Fig. 2.**
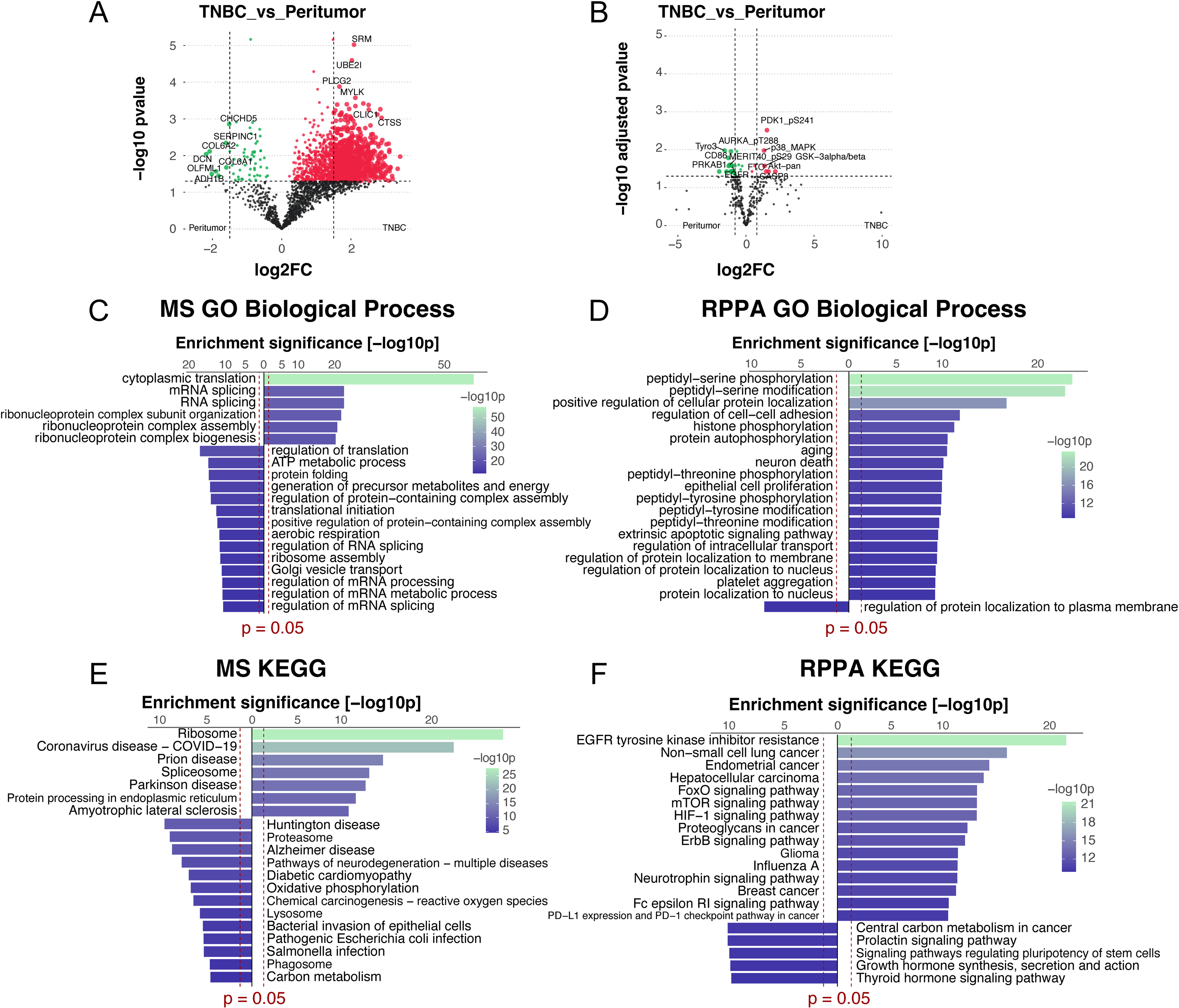
Differentially expressed proteins between TNBC and normal controls revealed by MS or RPPA. A/B. Volcano plots showing differentially expressed targets between TNBC and normal tissue for MS and RPPA, respectively. Log_2_FCs are set to 1.5 and 0.8 respectively. P values, either adjusted (for MS) or non-adjusted (for RPPA) were set to 0.05 (dashed lines). C/D. GO-BP enrichment of differentially expressed proteins from MS or RPPA (both log2FC and p value cutoffs were applied). Enrichment p values (cutoff=0.05) were shown by red dashed lines. E/F. KEGG pathway enrichment of differentially expressed proteins from MS or RPPA (both log2FC and p value cutoffs were applied). Enrichment p values (cutoff=0.05) were shown by red dashed lines.

Gene ontology-based pathway enrichment analysis revealed distinct regulatory patterns between MS and RPPA. MS highlighted processes such as RNA splicing, translation regulation, and metabolic processes, while RPPA focused more on post-translational modification and cellular functions such as proliferation, adhesion, and apoptosis (Fig. 2C/D). KEGG pathway analysis further underscored the individual regulatory patterns, with MS highlighting regulation in neurodegeneration, oxidative processes, and RNA/protein synthesis. RPPA demonstrated dynamic regulation within multiple cancers and key oncogenic signaling pathways, including EGFR, HER2, mTOR, FoxO, HIF-1, and PD-1/DP-L1 (Fig. 2E/F).

### Overlapping proteins identified by MS and RPPA

We next investigated the overlaps of proteins profiled by MS and RPPA. Since MS in this study did not identify protein phosphorylation and other post-translational modifications (PTM), only unique total proteins identified from both methods were compared. A total of 61 overlapping proteins were identified (Supplementary S7), which account for 2.4% of the 2,583 proteins from MS and 27.6% of 221 unique total proteins from RPPA (Fig. 3A). Among all DE proteins, 8 overlapping proteins were identified, accounting for 1% MS DE proteins and 13% of RPPA DE proteins (Fig. 3B). This partially explains the low overlap in pathways derived from GO and KEGG enrichment analyses, wherein only 7 and 17 pathways overlapped in either enrichment profiling (Fig. 3C/D).

**Fig. 3.**
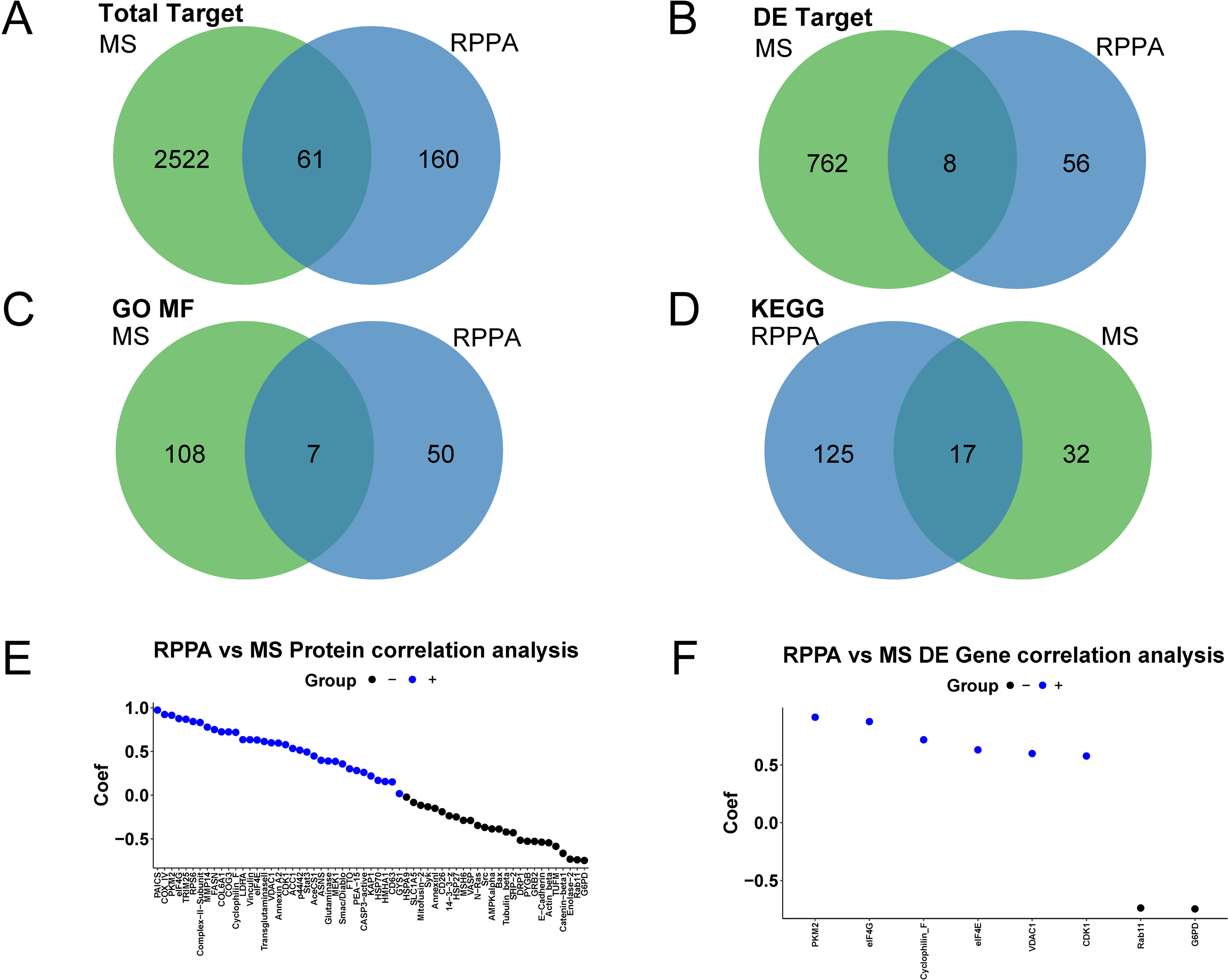
Overlapping proteins between MS and RPPA. A/B. Venn diagrams showing overlaps of all or differentially expressed proteins between TNBC and normal tissue profiled by MS and RPPA. C/D. Venn diagrams showing matching GO terms (MF) and KEGG pathways enriched in MS and RPPA profiling, respectively. E. Ranked matching targets correlation between MS and RPPA across samples. Blue dots show positively correlated proteins and black dots show negatively correlated proteins. F. Ranked correlation of matched differentially expressed proteins between MS and RPPA across samples. Blue dots designate positively correlated proteins and black dots designate negatively correlated proteins.

As both MS and RPPA provided quantitative proteomic data, we then analyzed the correlation between their matched targets. Of the total 61 unique proteins matched, a dynamic range of quantitative correlation was observed across shared targets. Genes such as PAICS, COX4l1, PKM had the highest correlation (R^2^>0.9), while other signaling regulators, including PRS6, MAPK3, STAT3 had moderate positive correlation (Fig. 3E). Approximately 55% of proteins showed a weak to strong positive correlation (Fig.3E). A further comparison focusing on the 8 matched DE proteins showed an overall 75% (6 out of 8) relatively-strong positive correlation (Fig. 3F).

### RPPA reveals the key pathway activation in TBNC

We then analyzed the proteomic data for EGFR and ERBB2 signaling, two critical drivers in TNBC. From our RPPA data, TNBC tumors, in comparison to paired normal samples, exhibited lower EGFR total protein levels (p<0.05) with lower phosphorylation levels at tyrosine residues 1068 (Fig. 4A). In consistent, a canonical downstream PI3K/AKT signaling was also downregulated in TNBC compared with peritumor tissues, shown by lower phosphorylation of AKT at residues threonine 308 and serine 473, the 2 major activating phosphorylation sites of the kinase, indicating that AKT was not activated in TNBC, despite elevated total AKT levels (Fig. 4B). AKT inactivation in TNBC was further demonstrated by downregulation of GSK3α/β phosphorylation, a downstream substrate of the AKT kinase (Fig. 4C). Our results are consistent with the public TCGA RPPA data of the TNBC samples, which also show EGFR phosphorylation at Tyr1068 is significantly downregulated, together with lower levels of AKT phosphorylation at both Thr308 and Ser473 (Supplementary Fig. 1).

**Fig. 4.**
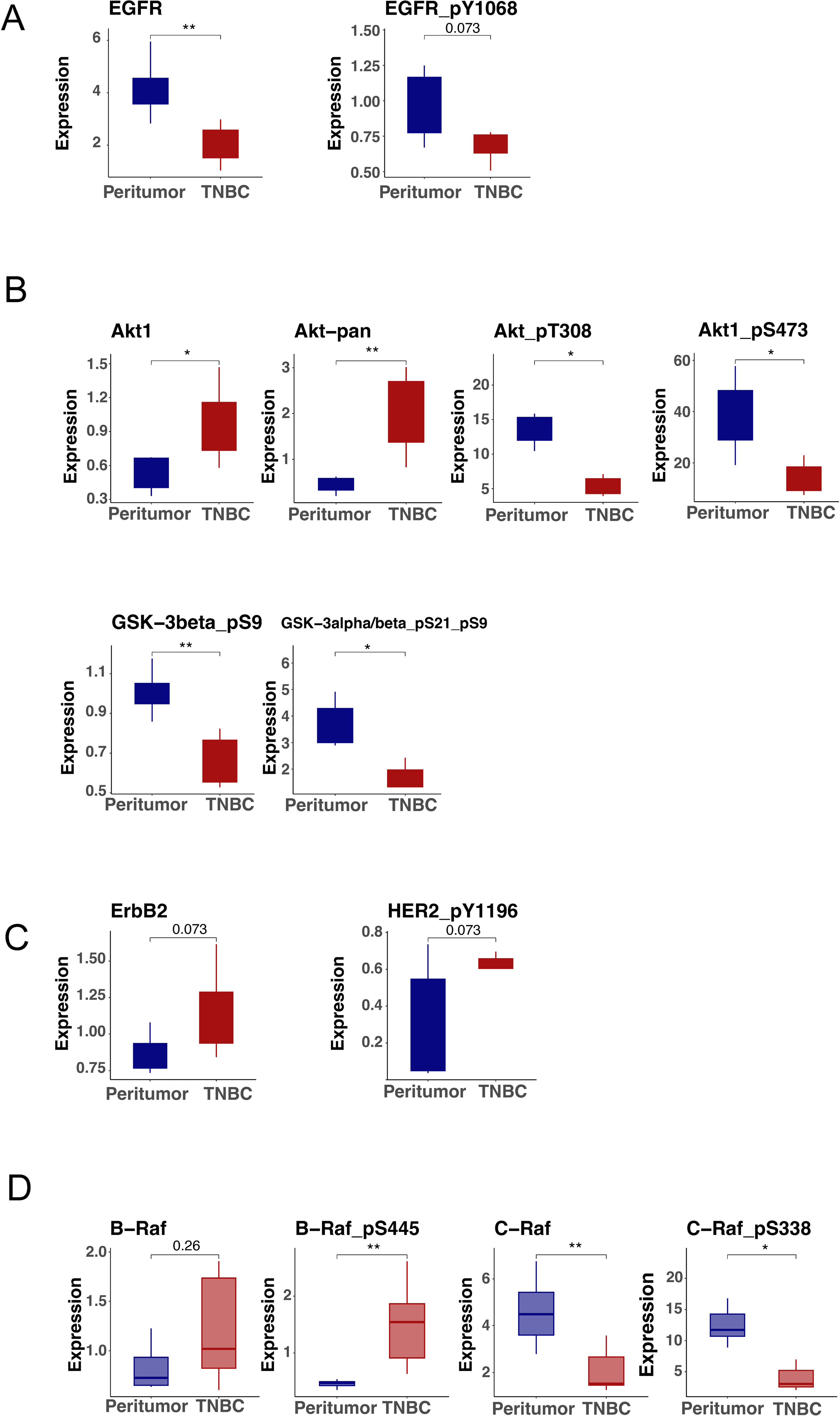
EGFR downregulation and ERBB2 upregulation in TNBC revealed by RPPA. A. EGFR total protein and pY1068 between TNBC and normal controls (p values). B. AKT1, pan-AKT, AKT pT308 and pS473, GSK-3β pS9 and GSK-3α/β pS21/S9 between TNBC and normal controls. C. HER2, HER2 pY1196 between TNBC and normal controls. D. B-Raf, B-Raf pS445, C-Raf and C-Raf pS338 between TNBC and normal controls. All comparisons were carried out using Wilcoxon ranked sum test with either p-values shown or asterisks representing p<0.05 (*), p<0.005 (**) and p<0.0005 (***), respectively.

Notably, our RPPA data showed that ERBB2 signaling was activated in TNBC as compared to their paired normal controls, which was indicated by increased HER2 total protein and its phosphorylation on tyrosine residues 1196 (Fig. 4C). HER2 activation triggers the downstream Raf/MEK/ERK signaling cascade. Our RPPA data further showed significantly increased B-Raf phosphorylation at serine residue 445 (p<0.05) as a likely activation axis through HER2 activation, but not C-Raf (Fig. 4D).

In contrast, no protein phosphorylation, which usually reflects protein activation status, can be conveniently captured in MS. Only changes of total proteins can be used to evaluate activation of signaling transductions. To compare the capacities of MS and RPPA in revealing signal transductions in TNBC, we analyzed changes of proteins identified in our MS and RPPA in KEGG pathways closely related to TNBC including EGFR, ERBB, PI3K/AKT, and mTOR pathways. As expected, while RPPA identified fewer proteins than MS, it revealed significantly more proteins that change in a variety of signal pathways (Supplementary Fig. 2A, B, C, D). Taking the EGFR pathway as an example, RPPA identified 32 proteins in the pathway, while MS identified 14 proteins (Supplementary Fig. 2A). The results showed that RPPA, designed for phosphoproteins and low-expression signaling proteins especially in cancer signal transductions, offers a more detailed insight into signaling pathway changes than MS. As two complementary proteomic technologies, the targeted proteomic RPPA excels in profiling signaling networks in diseases like cancer, while the de novo MS technology excels in revealing previously unknown mechanisms.

### Parallel MS and RPPA proteomic profiling of 293T cells under basal and FBS-stimulated conditions

Since MS and RPPA are widely used on cell cultures in addition to tissue samples, we further assessed the outcomes and performance of these two technologies on cell line models. We first tested on 293T cells under serum starvation and FBS stimulation conditions, representing the same cells under different physiological conditions. Using the same DDA data acquisition mode in MS, we obtained 2,317 proteins from 293T cell samples. Similarly, 305 proteins including 230 total proteins and 75 phosphoproteins were obtained by RPPA from the same samples. Hierarchical clustering based on both profiling exhibited clear separations between starvation and stimulation groups (Fig. 5A, Supplementary S8).

**Fig. 5.**
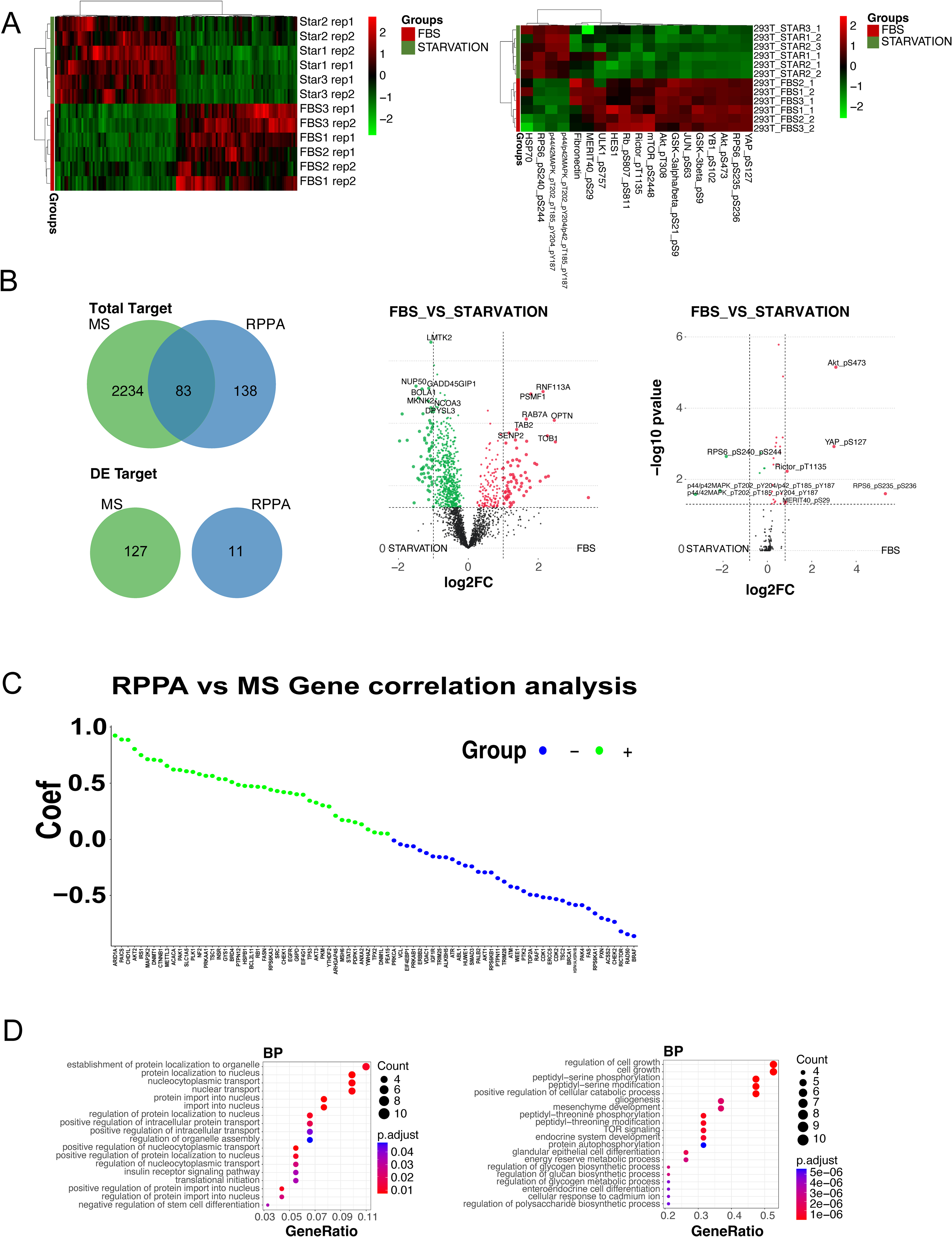
Parallel MS and RPPA profiling on 293T cells under basal and stimulated conditions. A. Hierarchical clustering of differentially expressed proteins/modified proteins for MS (left) and RPPA (right) of 293T cells under basal or FBS-stimulated conditions. Experiments were done in triplicates. B. Volcano plots showing differentially expressed proteins or modified forms between two treatment conditions. For MS and RPPA, Log_2_FCs are set to 1.5 and 0.8 respectively. Right panel shows unique and shared targets between MS and RPPA for all identified proteins (upper) or differentially expressed proteins (lower). C. Target-wise correlations across samples of shared proteins (presented as gene names). D. GO enrichment based on differential expressed proteins profiled through MS (left) or RPPA (right) respectively. P-values (adjusted) and counts are illustrated on the right side.

We compared the unique total proteins identified by MS and RPPA, revealing 83 overlapping proteins, which constitute 3.6% (83/2317) of MS proteins and 37.6% (83/221) of RPPA proteins (Fig. 5B). The overlapping rates are higher than those in the TNBC tissues (2.4% and 27.6%), likely due to the analysis of the same cells. A total of 127 differentially expressed (DE) proteins were identified from MS, and 11 from RPPA. Interestingly, no overlap was observed among the DE proteins identified by MS and RPPA (Fig. 5B). Only about 50% of the overlapping proteins showed positive correlation (Fig. 5C). Using GO enrichment analysis, we identified distinct biological patterns associated with each technology. In MS, significantly altered proteins were prominently enriched in pathways related to protein transportation and shuttling in response to FBS stimuli (Fig. 5D, left). In contrast, RPPA, which is designed to focus on intra-cellular signaling, exhibited most altered proteins enriched in pathways associated with cell growth regulation, serine/threonine phosphorylation, TOR signaling, as well as other cellular differentiation and development processes (Fig. 5D, right).

### Parallel MS and RPPA proteomic profiling of different cell lines

We performed MS with DDA for TNBC and 293 cells to compare with parallel RPPA. Recognizing the increasing applications of MS under the DIA scanning model, which is expected to yield more proteins than DDA with more accurate quantification, we further performed MS with DIA and RPPA on two different cell lines (gastric cancer cells MKN7 and ovarian cancer cells OVCAR3). In MS with DIA, around 6,000 proteins were successfully identified and quantified, a significantly larger number compared to the proteins identified in TNBC tissue or 293T cells using MS with DDA. The same 305 proteins were identified by RPPA in the same samples. Both MS and RPPA profiling effectively differentiated between the two cell lines (Fig. 6A, Supplementary S9).

**Fig. 6.**
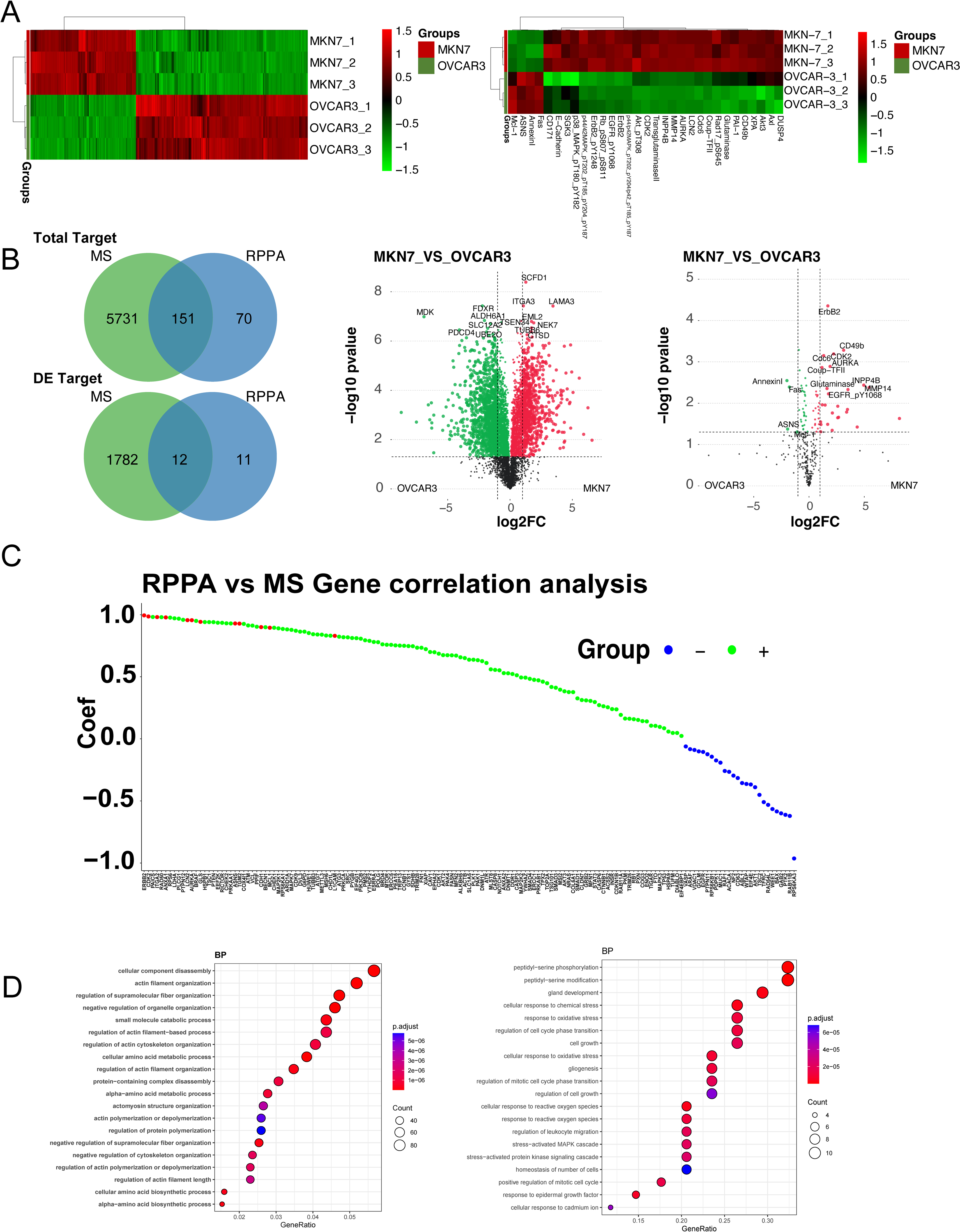
Parallel MS and RPPA profiling on different cell lines. A. Hierarchical clustering of differentially expressed proteins/modified proteins for MS (left) and RPPA (right) between different cell lines (MKN-7 and OVCAR-3). Experiments were done in triplicates. B. The left panel shows unique and shared targets between MS and RPPA for all identified proteins (upper) or differentially expressed proteins (lower). Corresponding volcano plots showing differentially expressed proteins or modified forms between two cell lines. For MS and RPPA, Log_2_FCs are set to 1.5 and 0.8, respectively. C. Target-wise correlations across samples of shared proteins (presented as gene names) between MKN7 and OVCAR3. Positive correlation shown in green and negative correlation shown in blue. Shared differentially expressed targets shown in red. D. GO enrichment based on differential expressed proteins profiled through MS (left) or RPPA (right) respectively. P-values (adjusted) and counts are illustrated on the right side.

The overlapping unique proteins from MS and RPPA were 2.6% (MS:151/5882) and 68.3% (RPPA:151/221), respectively. For DE proteins (under cutoffs: logFC=1 and p-value=0.05 removing targets with missing values in replicates), the overlapping rates are 0.67% in MS (12/1794) and 52.2% in RPPA (12/23) (Fig. 6B). About 70% of overlapping proteins showed positive correlation (Fig. 6C), higher than that in 293T cells, likely due to a better qualification of DIA compared with DDA. Furthermore, all matched DE proteins (designated with red dots) exhibited positive correlation, confirming their quantitative consistency (Fig. 6C). Similarly, GO enrichment analysis showed distinct biological patterns associated with MS and RPPA (Fig. 6D).

Collectively, these data further support the overall feasibility of our method for profiling in vitro cell lines across various biological states, and highlight the technical robustness and complementary nature of the two proteomic technologies in profiling biological samples.

## Discussion

In this study, we employed two prevalent proteomic technologies, namely label-free LC-MS and RPPA, to establish an efficient and streamlined workflow for sample preparation to data analysis in both tissue and cell line settings. Label-free LC-MS, without requirement for specific LC-MS reagents and complex MS run setup, was assessed with both DDA or DIA to compare with RPPA in parallel. To prepare protein samples suitable for both technologies, we initially used detergent-rich and chaotropic compound-containing lysis buffer typically used in RPPA procedures to solubilize membrane and nuclear proteins. Subsequently, these protein samples underwent acetone precipitation, and the buffer was changed to meet the requirements of downstream LC-MS applications, with starting materials of 50 μg and a typical injection volume of 2 μg.

From the data analysis perspective, both proteomic technologies effectively captured the differences in protein profiles between sample groups (TNBC tumor vs. peritumor normal, different cell lines, and cells under different conditions). However, they diverged in their ability to uncover distinct underlying biological mechanisms, as revealed by the results of GO and KEGG analyses. Differential expression analysis highlighted a relatively low number of overlapping targets identified through both methodologies, suggesting inherent technical characteristics associated with each method.

As for LC-MS, our objective was to implement a straightforward yet rigorous run condition using a high-quality instrument, employing a single column with two technical replicates for each sample within an acceptable instrument time frame (1 hour). For tissue samples, using standard data analysis tools, 3,356 proteins were identified with 2642 being quantified with decent reproducibility, which was typical in a standard DDA LC-MS run. However, with the aid of longer gradients, tandem mass tags (TMT) or data-independent acquisition (DIA), reproducibility and detection capability may be further enhanced, as in the CPTAC analyses.

In the comparison with an external database, Our MS data displayed nearly no correlation with the CPTAC data for TNBC samples. This lack of correlation could be attributed not only to the limited sample number being compared (18 TNBC samples plus 3 normal samples from the CPTAC database) but also to the distinct experimental setup. The reference studies used TMT 10-plex quantification and pre-fractionation, with a total of 110-minute LC-MS/MS gradient runs. On the other side, TNBC samples profiled through RPPA exhibited relatively better correlation and this was not only due the larger sample size being compared but also our experimental setup that was similar to the method applied in TCGA breast cancer RPPA profiling. Nevertheless, the inter-tumor sample correlation was inferior to inter-normal correlation and this was potentially due to the heterogeneity of TNBC subtype or the genetic discrepancy across different racial background (East Asian/Chinese versus Caucasian/Africa American).

In both tissue and cell lines, the quantitative comparison of shared targets between LC-MS and RPPA demonstrated various degrees of quantitative correlation. This is consistent to a previous study showing an overall 60% positive correlation between LC-MS and RPPA data [32]. RPPA, functioning akin to a high-throughput immuno dot blot assay, is more sensitive in quantifying low-abundance expressors, rendering it more reproducible for dissecting signaling transductions [8]. In contrast, LC-MS involves more intricate pre-analytical steps that may impact quantitative accuracy by measuring peptide abundance, posing a challenge when dealing with complex sample types like tissues. Regardless of these facts, more than half of the matching targets showed consistent quantitative results ensuring the robustness of our data across two platforms.

Furthermore, we compared key targets in major oncogenic signaling pathways in TNBC. Our results showed down-regulation of EGFR protein and its activity compared to paired normal tissues. Previous studies utilizing either MS or RPPA for discovery proteomics demonstrated EGFR up-regulation in TNBC compared to luminal A/B and HER2+ breast cancer [19, 32]. However, whether the observed down-regulation of EGFR signaling compared with paired normal tissues is specific to the samples requires further investigation. If suitable anchoring samples become available, we could compare our RPPA data with external TCGA RPPA data for breast cancer samples using RBN-based normalization for a more comprehensive analysis [33]. RPPA profiling of the HER2 pathway revealed elevated expression and HER activity in tumor samples. This finding aligns partially with previously reported data where highly phosphorylated HER2 was found in HER2-negative breast cancer tissues and cell lines [34]. Our exploratory results also demonstrated that, in connection with HER2 activation, another canonical HER2 downstream pathway, RAF/MEK/ERK, may be triggered, primarily through B-Raf, the most potent Raf isoform.

Finally, the assessment of the two parallel proteomic profiling on cell line samples further validated the combined complementary technologies. In 293T cells, while MS captured global changes at downstream of signaling pathways following serum stimulation, RPPA was capable of detecting subtle and transient pathway alterations intracellularly that directly link with growth stimulation at early time points, highlighting its key feature in quantitative measurement of intracellular signal cascades.

In summary, we integrated and evaluated two complementary proteomic technologies using the same samples. The straightforward LC-MS, by avoiding complex sample preparation methods such as filter-aided sample preparation (FASP) or solid-phase-enhanced sample preparation (SP3), as well as pre-labeling strategies like TMT, enables a quick experimental turnaround time and is cost-efficient. With a LC run coupled with high-resolution MS using either DDA or DIA, thousands of proteins can be quantified. When coupled with RPPA, additional lower-abundance proteins and corresponding PTMs can be obtained, providing a deeper insight into signaling transduction. We demonstrated the advantages of incorporating LC-MS and RPPA as two complementary discovery proteomics technologies, allowing for a complimentary proteomic profiling of both tissue and cell line samples, providing comprehensive mechanistic insights.

## Acknowledgments

We thank other team members at Fynn Biotechnologies for conducting RPPA and related work and members at Department of Chemistry, Tsinghua University to optimize the LC-MS workflow.

## Data availability statement

All LC-MS and RPPA raw data were deposited at GitHub. This article contains Supplementary Table 1, Supplementary Figures, and Supplementary files S1-9 in excel spreadsheet format.

## Author contributions

Nan Wang: Methodology, Formal analysis, Writing-Original Draft, Project coordination; Yiying Zhu: Methodology, Experimentation, Writing-Review & Editing; Lianshui Wang: Data analysis; Wenshuang Dai: Software, Formal analysis, Data curation and visualization; Taobo Hu: Sample acquisition, Supervision, Review & Editing, Zhentao Song: Software, Formal analysis, Data curation; Xia Li: Supervision, Experimentation; Qi Zhang: Experimentation; Jianfei Ma: Experimentation; Qianghua Xia: Data analysis; Jin Li: Data analysis; Mengping Long: Resources, Supervision, Writing-Review & Editing, Zhiyong Ding: Conceptualization, Methodology, Review & Editing, Supervision.

## Abbreviations

ACN: acetonitrile
AMBIC: ammonium bicarbonate
BCA: bicinchoninic acid assay
CF: correction factor
CPTAC: clinical proteomics tumor analysis consortium
DDA: data-dependent acquisition
DIA: data-dependent acquisition
DIGE: difference in gel electrophoresis
EGFR: epidermal growth factor receptor
EGTA: ethylene glycol tetraacetic acid
ESI: electrospray ionization
FA: formic acid
FASP: filter aided sample preparation
FF: fresh frozen
GDC: genomic data common
GO: gene ontology
HPLC: high-performance liquid chromatography
KEGG: Kyoto Encyclopedia of Genes and Genomes
LC-MS: liquid chromatography-mass spectrometry
MALDI: matrix absorbed laser dissociation ionization
MRM: multiple reaction monitoring
PTM: post-translational modification
RPPA: reverse phase protein arrays
SDS: Sodium dodecyl sulfate
SP3: solid phase enhanced sample preparation
TCGA: the cancer genome atlas
TFA: trifluoroacetic acid
TMT: tandem mass tag
TNBC: triple-negative breast cancer

