## Supplementary Figures for "Parallel Analyses by Mass Spectrometry (MS) and Reverse Phase Protein Array (RPPA) Reveal Complementary Proteomic Profiles in Triple-Negative Breast Cancer (TNBC) Patient Tissues and Cell Cultures"

### Figure S1

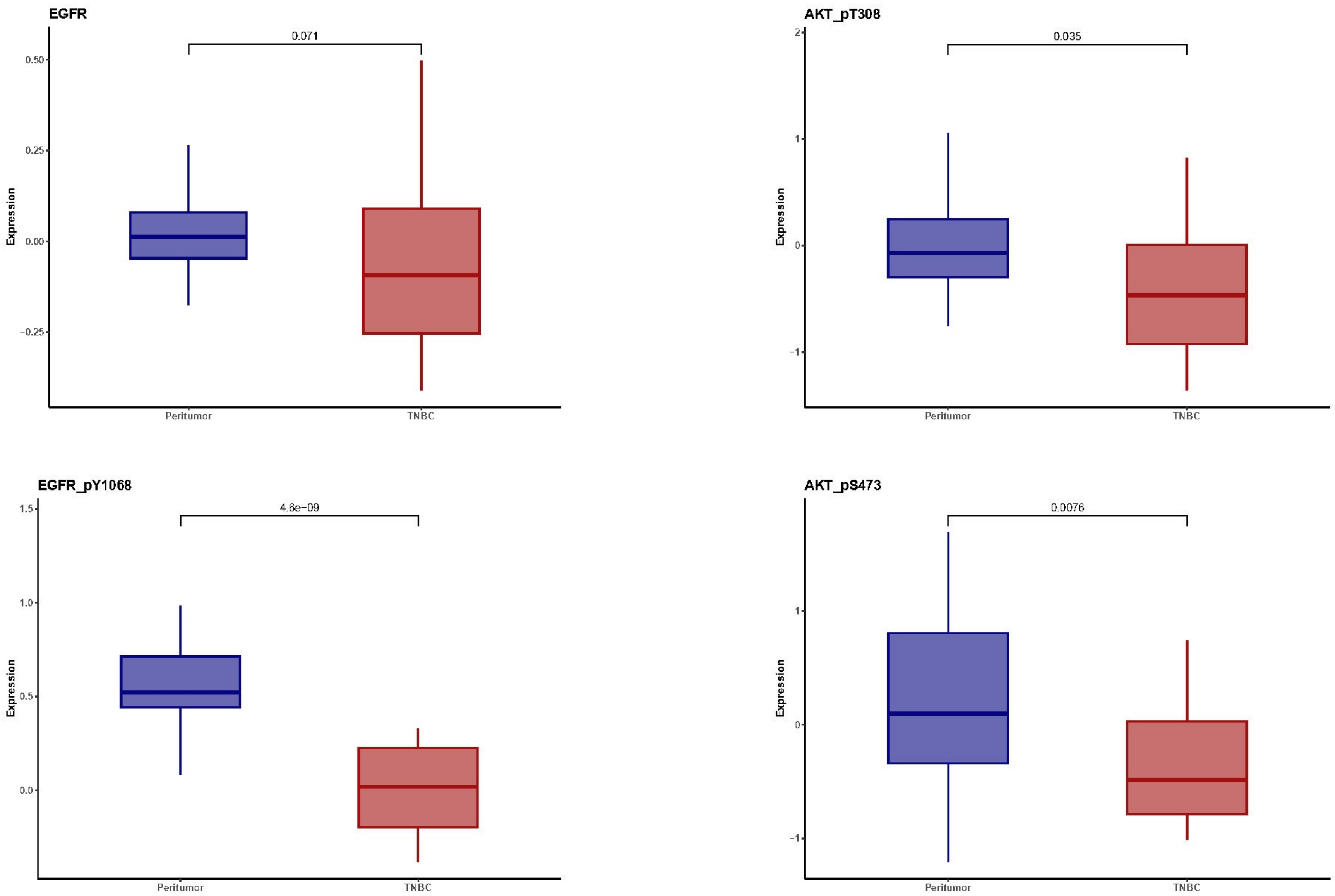
EGFR
0.071
0.50
0.25
0.00
−0.25
Expression
Peritumor
TNBC
AKT_pT308
0.035
2
1
0
−1
Expression
Peritumor
TNBC
EGFR_pY1068
4.6e−09
1.5
1.0
0.5
0.0
Expression
Peritumor
TNBC
AKT_pS473
0.0076
1
0
−1
Expression
Peritumor
TNBC

### Figure S2 A

MS

RPPA

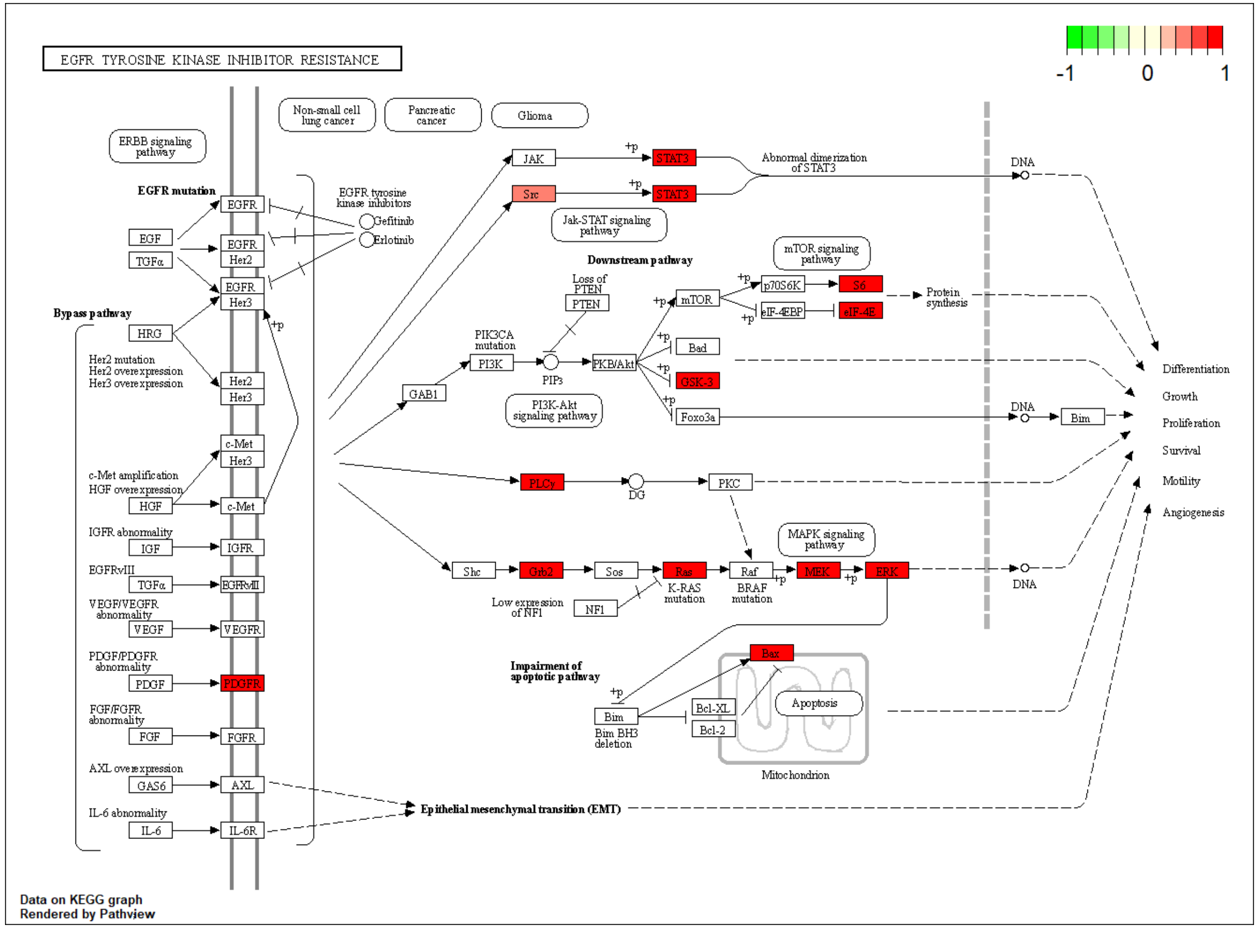


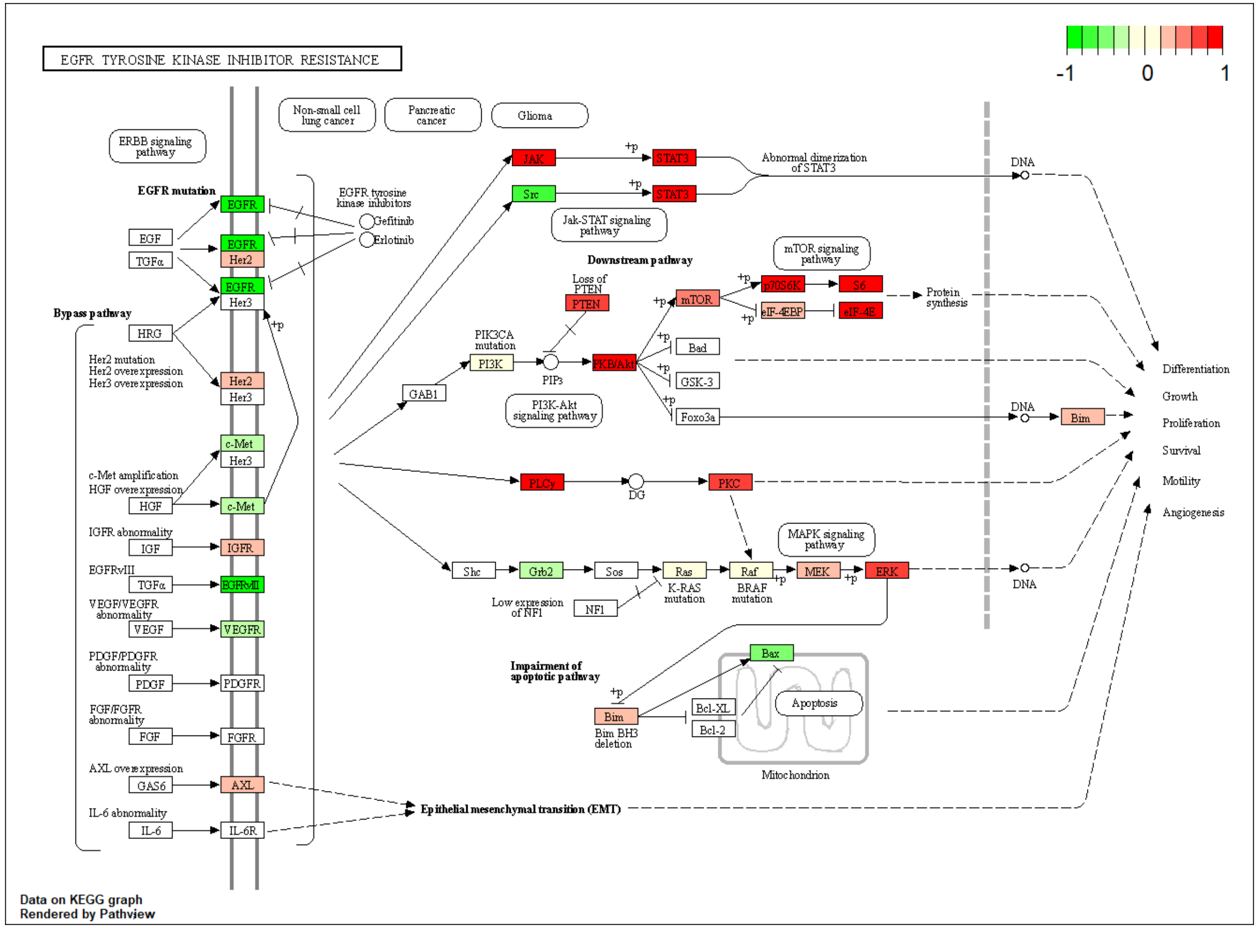


TNBC VS Peritumor

### Figure S2 B

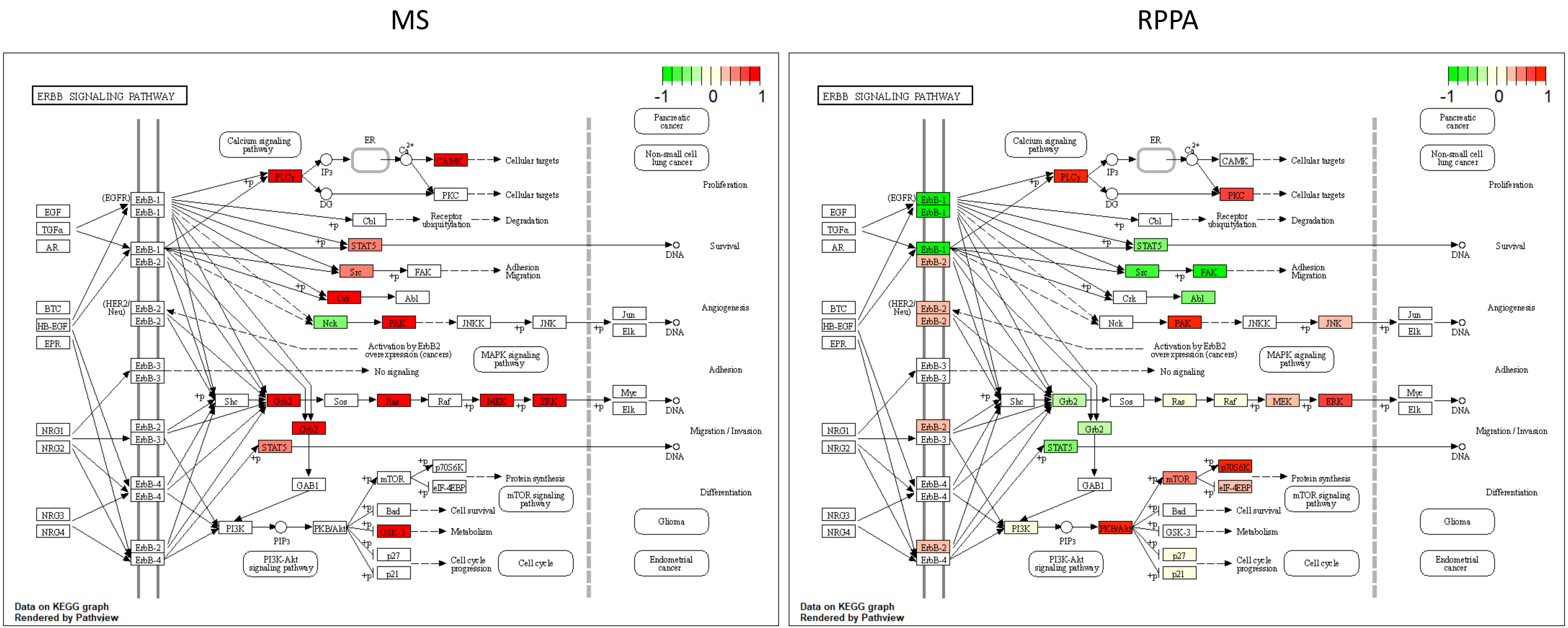


TNBC VS Peritumor

### Figure S2 C

MS

RPPA

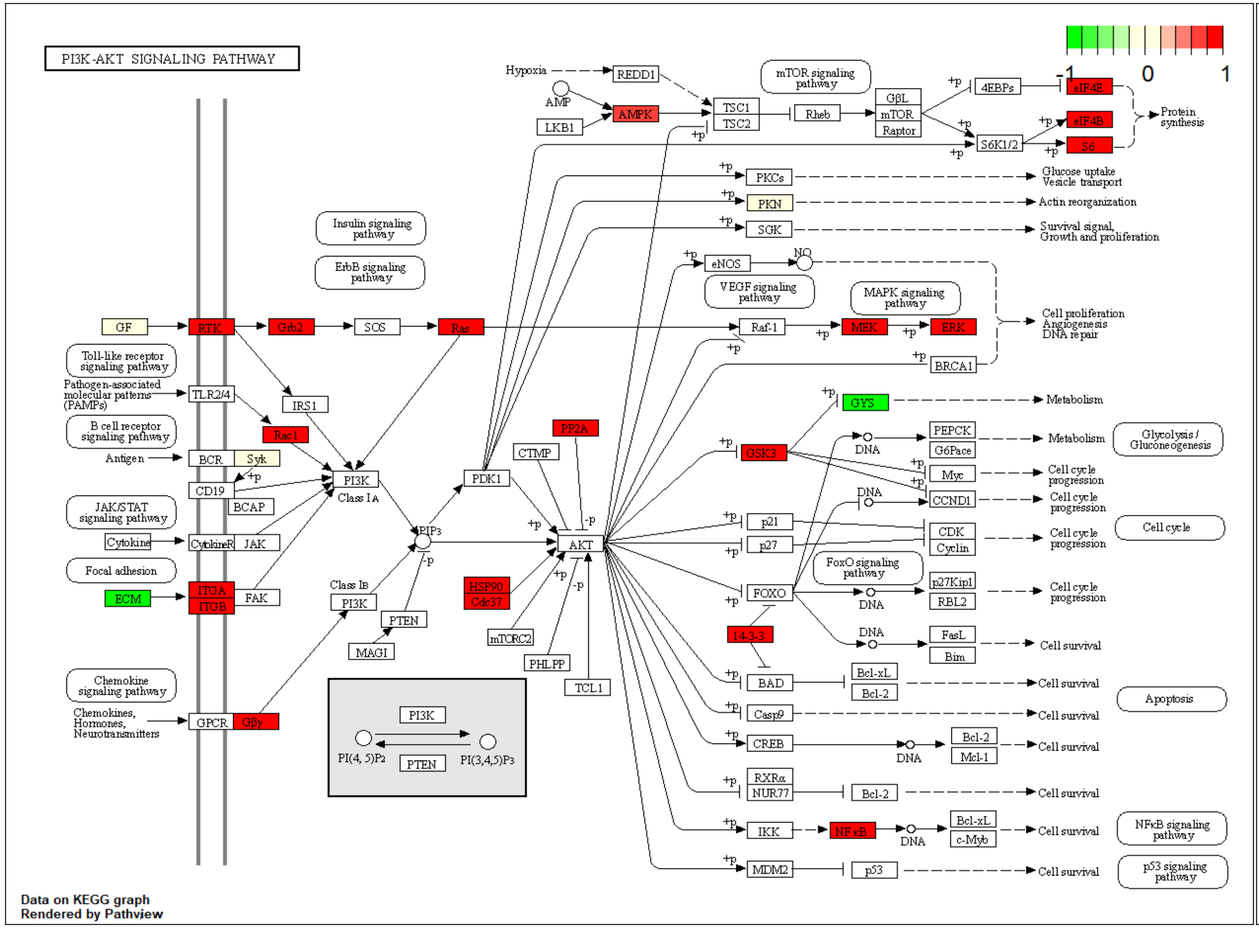


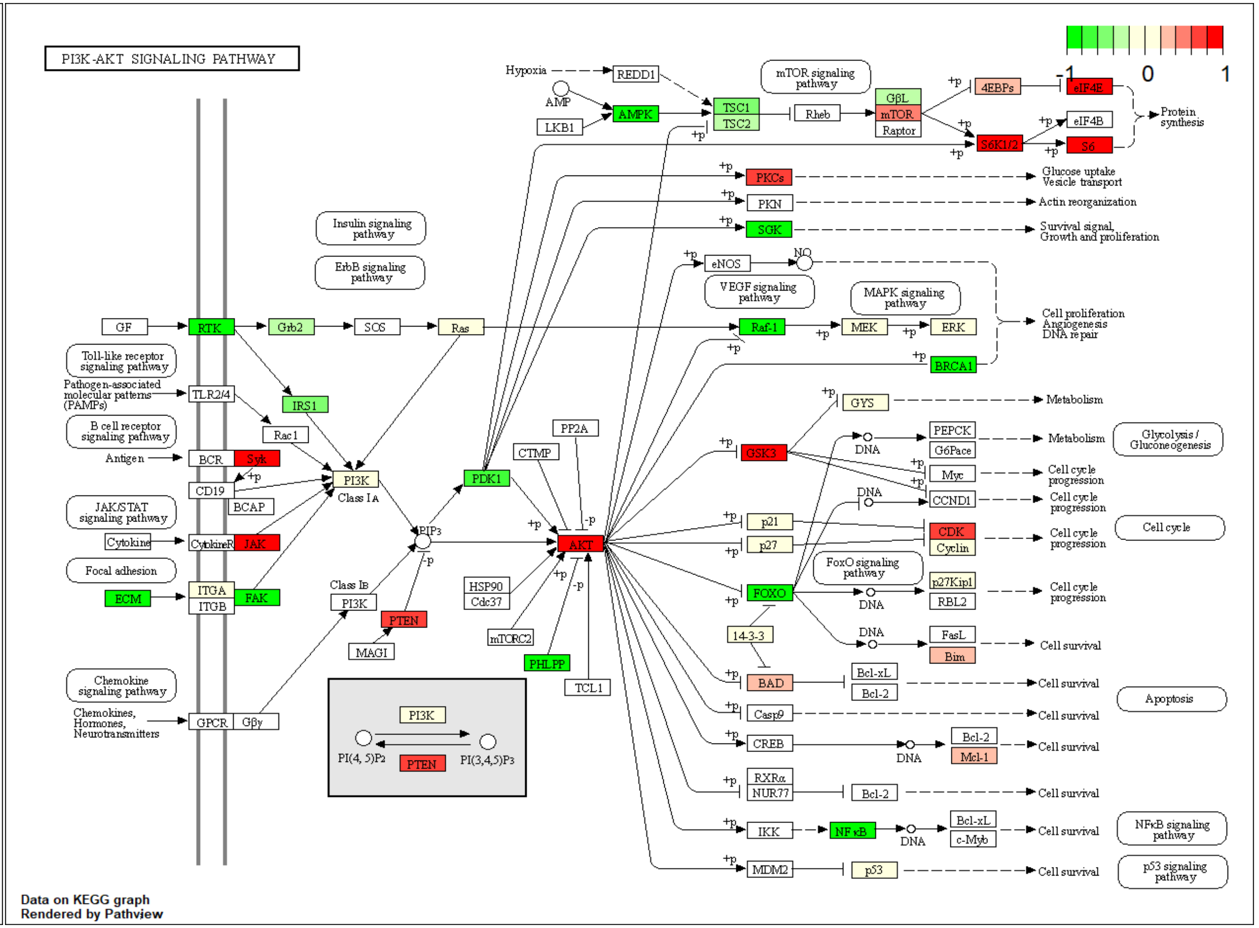


TNBC VS Peritumor

### Figure S2 D

MS

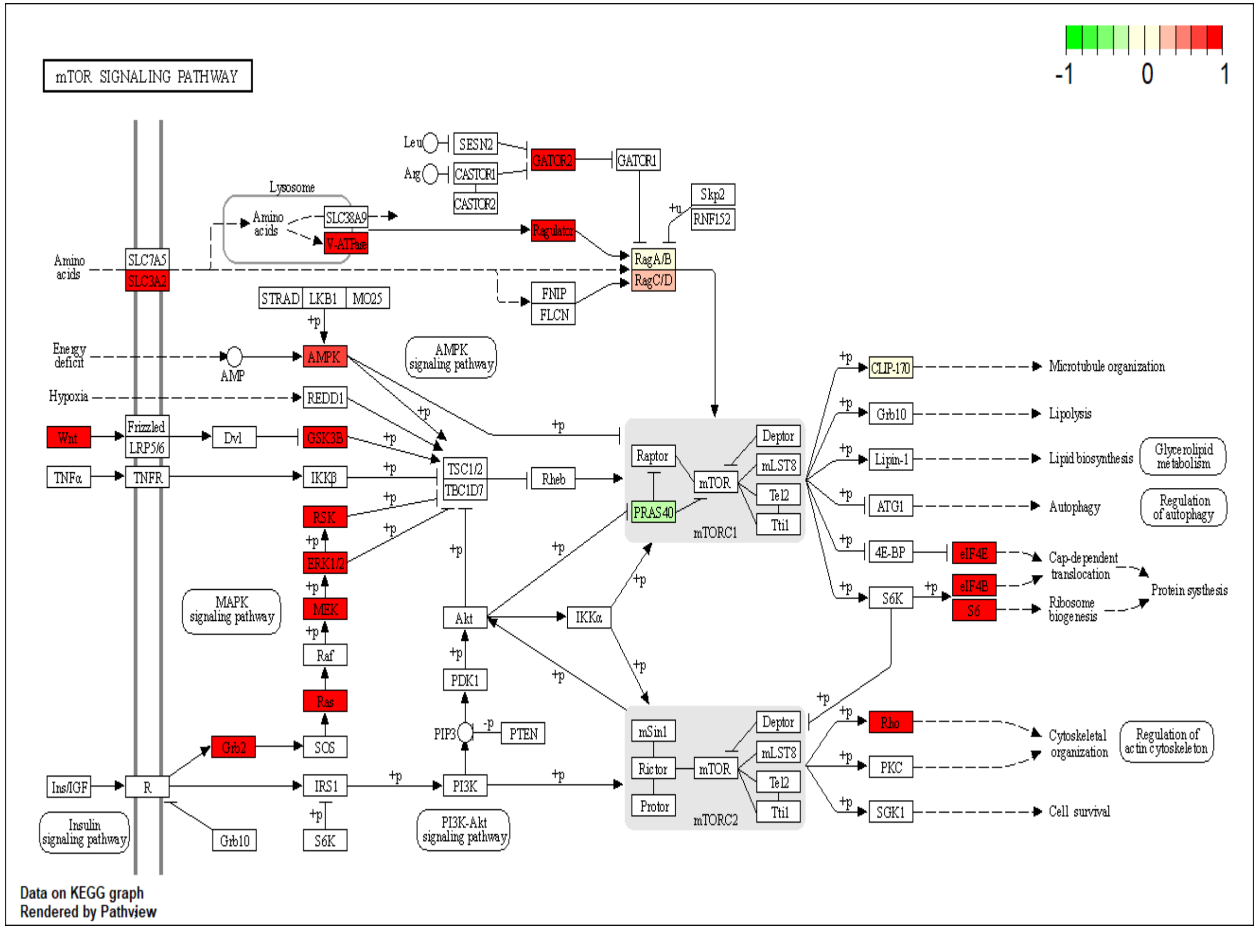


RPPA

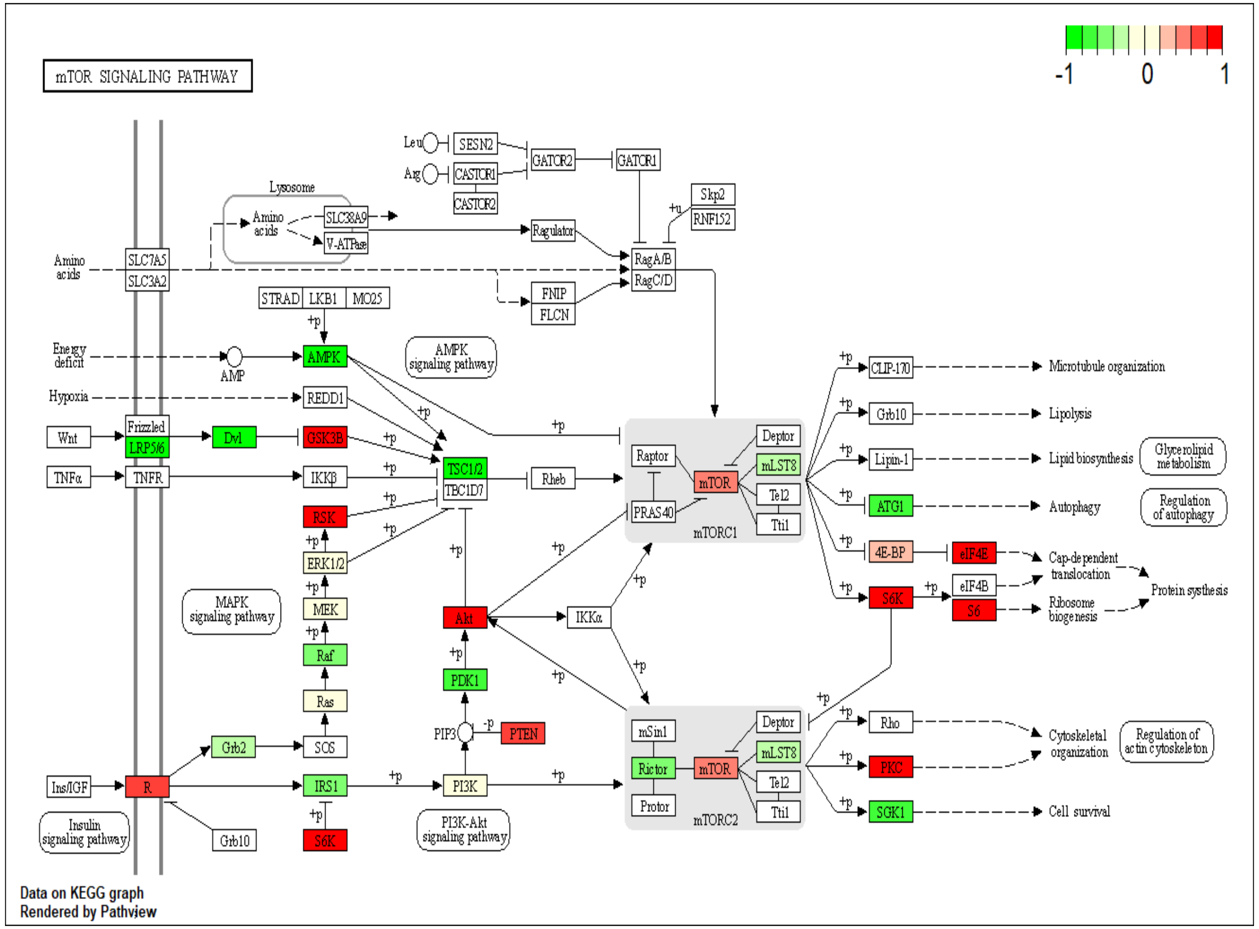


TNBC VS Peritumor
