## Supplementary Figure Legends for "Parallel Analyses by Mass Spectrometry (MS) and Reverse Phase Protein Array (RPPA) Reveal Complementary Proteomic Profiles in Triple-Negative Breast Cancer (TNBC) Patient Tissues and Cell Cultures"

Fig. S1. EGFR signaling downregulation in the TCGA TNBC samples revealed by the public RPPA database. Total EGFR, EGFR pY1068, AKT pT308 and AKT pS473 between TNBC and normal controls. All comparisons were carried out using Wilcoxon ranked sum test with p-values shown.

Fig. S2 A~D. KEGG pathways of EGFR, ERBB, PI3K/AKT, and mTOR. Based on differential expression analysis, KEGG Pathway Visualization was done with "pathview" package (R package). The value of log2FC (Fold Change is the logarithm base 2) of the comparison between TNBC and normal controls is color-annotated. The log2FC values of individual targets are represented by colors (Red-green transition).
